# COMPARATIVE RADIOPATHOLOGY OF MALE REPRODUCTIVE ORGANS IN SPRAGUE DAWLEY RATS ADMINISTERED WITH *Curcuma longa* AND *Ocimum sanctum* AGAINST ACUTE GAMMA IRRADIATION

**DOI:** 10.1101/2022.12.02.518900

**Authors:** Meikunu Thapru, Anirudh A Nair, Rakibul Hoque, Mousumi Namasudra, Arun Kumar De, Saktipada Pradhan, Rabindra Nath Hansda, S. K. Mukhopadhayay, Samiran Mondal

## Abstract

**Objective:** The study was designed to evaluate the deleterious e□ects of acute lethal gamma irradiation on male reproductive organs of Sprague Dawley rats and the radioprotective potential by *Curcuma longa* and *Ocimum sanctum* in experimental set up.

**Methods:** 25 Male Spraque Dawley rats were taken and randomized into 5 groups (T1, T2, T3, T4, T5). Each group containing 5 animals. The groups were named as Group T1 (Negative Control), Group T2 (irradiation positive control), Group T3 (irradiation and *Curcuma longa*), Group T4 (irradiation and *Ocimum sanctum*), Group T5 (irradiation, *Curcuma longa* and *Ocimum sanctum*). The rats in groups T1 and T2 were given a daily dosage of normal saline for 14 days. Group T3 and T4 rats received aqueous extracts of *Curcuma longa* (40 mg/kg) and *Ocimum sanctum* (40 mg/kg), respectively, whereas group T5 rats received a combination of aqueous extracts of *Curcuma longa* (40 mg/kg) and *Ocimum sanctum* (40 mg/kg) for 14 days. prior to gamma irradiation. Animals of group T2, T3, T4, T5 were irradiated with 10 Gy of γ – radiation on 15th day.

**Results:** Exposure of animals to 10 Gy gamma radiation resulted in significant decrease in body weight and testicular weight. Histopathological observations revealed testicular degenerative changes in the seminiferous tubules with primary spermatocytes being most affected and epididymis showed decreased spermatids density with cell debris in the lumen. Pretreatment with *Curcuma longa* at 40mg/kg for 14 days alleviated the radiation induced pathological changes in testis and epididymis. *Curcuma longa* demonstrated much better protective effect in combination with *Ocimum sanctum* in alleviating the radio pathological changes showing the possibility of synergistic effect. However, administration of *Ocimum sanctum* alone at 40mg/kg for 14 days did not reveal any significant radioprotective property after histomorphological evaluation.

**Conclusion:** Gamma irradiation of 10 Gy induces severe radio pathological damage to organs of male reproductive system of Spraque Dawley rats. Pre-treatment with *Curcuma longa* prior to radiotherapy may be effective in preventing radiation-induced damage in the male reproductive organs. Better radioprotection can be achieved with the combination of *Curcuma longa* and *Ocimum sanctum* than using alone.

## 1. INTRODUCTION

Ionizing radiations have a significant influence on living cells, and the rapid growth of technology has increased human exposure to ionising radiations.

Ionizing radiation is used in medical diagnostics, radiotherapy, and sterilization (Geri *et al*., 2019). Despite advancements in clinical radiation treatment planning and delivery technologies, radiotherapy still has a high level of damage to normal tissues and organs (Bhandari, 2013). The effect of ionising radiations on living tissue is influenced by a variety of characteristics, including the total length of exposure, the source of radiation, the distance from the source, and the physical and metabolic properties of the living tissue. Radiation’s effects are regulated by both the dose and the duration of exposure (Haines *et al*., 2002).

Although all biological entities are vulnerable to ionising radiation, mammalian testes are far more sensitive. In both humans and animals, the testes are positioned outside the body and are susceptible to radiation damage. The dose and length of exposure to artificial radiation or treatment are directly proportional to testicular damage (Abuelhija *et al*., 2013).

The most visible effect of radiations on living tissue is the production of reactive oxygen species (ROS) and free radicals in exposed tissue (Tominaga *et al*., 2004). By producing reactive oxygen species (ROS), gamma irradiation promotes oxidative stress in testicular tissue, causing spermatogonia, spermatocytes, and spermatozoa to apoptose (Meistrich, 2013). Oxidative stress plays an important role in the pathogenesis of male infertility. Since, spermatogenic cells are constantly in mitosis or meiosis, they are especially vulnerable to radiation-induced reactive oxygen species (ROS). As a result, the testes are a highly radiosensitive organ with a diverse array of radiosensitive germ cells (Agarwal *et al*., 2006).

Radiation is most damaging to actively dividing cells or cells that are not fully mature, in a nutshell. Radiation damage is more sensitive in cells with a fast division and metabolic rate, as well as non-specialized and well-nourished cells. (Lehnert, 1975).

The effect of gamma ray on testicular organs of rats resulted in a reduction in the value of spermatogenic cells (Eissa *et al*., 2007). High doses of radiation can be lethal. When germ cells are exposed to radiation for a short period of time, they are killed and damaged, which has an impact on fertility (Wallach *et al*., 1986). Whether irradiation was given chronically at a low dose rate or abruptly at various total doses, the type and number of cells killed in the testis differed. Various germ cell connections in the seminiferous epithelium were altered over time as the maturation depletion process progressed (Dym and Clermont, 1970).

Various elements from natural and artificial sources have been examined and assessed in recent years in order to develop effective radioprotectors (El-Desouky *et al*., 2014). Radioprotective agents, also known as radioprotectors, are chemical or biological molecules are used to shield tissues and cells from the effects of radiation (Yi *et al*., 2018). Radioprotective compounds were employed as key components to limit caused harm, such as radiation-related mortality, before being exposed to gamma radiation. (Satoh *et al*., 2003). Radioprotective compounds are particularly valuable because, in addition to protecting normal tissue from radiation injury, they allow for higher doses of radiation to improve cancer control and possibly cure (Hosseinimehr, 2007); (Baliga *et al*., 2010). Radioprotective drugs, in particular, prevent the creation of reactive compounds (free radical scavengers), detoxify free radicals produced by radiation, target the stabilisation of important biomolecules, and stimulate the repair and recovery processes. (Mun *et al*., 2018).

Crude plant extracts have been shown to have radioprotective activity and significantly reduce cellular damage, including LPO and protein oxidation, among other types of degradation (El-Desouky *et al*., 2014) *Curcuma longa* has been found to protect normal organs from the effects of chemotherapy and radiotherapy, as well as serving as a chemo- and radiosensitizer for cancers in some cases (Jagetia, 2007). Curcumin have anti-inflammatory, phytonutrient, and bioprotective effects (Hossain *et al*., 2009). Curcumin is a potent antioxidant and one of the most effective free radical destroyers. (Chandra *et al*., 2007; Azza *et al*., 2011) *Ocimum sanctum* has anti-carcinogenic and radioprotective properties (Pattanayak *et al*., 2010). The leaves of *Ocimum sanctum* exhibit selective radioprotective capabilities at non-toxic quantities. *Ocimum sanctum* has the ability to shield the body’s DNA from harmful radiation (Uma Devi, 2006; Singh, 2005; Panda and Kar,1998; Devi and Gonasoundari, 1995).

## 2. MATERIALS AND METHODS

### 2.1. Approval from IAEC

This study was planned with a prior approval of Institutional Animal Ethics committee (IAEC) which is under supervision of CPCSEA, Government of India.

IAEC/WBLDCL/3/PROJECT/2021, dated 09/09/21 is the study approval number.

### 2.2. Experimental animal

25 Male Sprague Dawley were purchased from CPCSEA registered breeder; State Centre for Laboratory Animal Breeding, West Bengal Livestock Development Corporation Ltd, Kalyani, West Bengal. (CPCSEA registration number: 763/GO/RE/SL/03/CPCSEA). They were placed at animal house facility attached to the Department of Veterinary Pathology, WBUAFS in open cage system facility in controlled environment. The animals were acclimatized for a period of 5 days. Body weights were recorded at the start and end of acclimatization period. Animals were observed for mortality and morbidity twice daily. Grouping of animals was performed on the last day of acclimatization by body weight randomization. Each animal was identified by tail marking. Temperature and relative humidity were maintained at 20-26 °C and 30-70% respectively. Sterilized corn cob was used as bedding materials. The cages were washed with detergent and sanitized with absolute alcohol and hydrogen peroxide. Feed (rodent pellet feed) and water was supplied ad libitum. The maximum and minimum air temperature recorded during the period were 34^0^C and 26^0^C respectively. The average relative humidity ranged from 50% to 95%.

### 2.3. Chemical’s procurement

Extracts of *Ocimum sanctum* (Tulsi) and *Curcuma longa* (Turmeric) were procured from Just Jaivik, Ahmedabad, Gujarat and Asitis Nutrition, Bangalore, Karnataka, respectively.

### 2.4. Formulation

#### 2.4.1. *Curcuma Longa* extract (Turmeric) and *Ocimum Sanctum* extract (Tulsi) preparation

Dose of Turmeric and Tulsi was @40 mg/kg each, maximum oral dose volume was @ 10ml/kg, dose concentration was 10 mg/ml each. Every day, fresh formulation of extract was prepared for oral administration by adding sterile potable water to it and fed through oral gavage using sterilized oral metal gavage of 18G directly to stomach for 14 days. (Bhartiya *et al*., 2010).

### 2.5. Experimental design

2.5. Experimental design:
| Route of Administration | Oral gavage |  |  |  |  |
| --- | --- | --- | --- | --- | --- |
| No of Animals/Group | 5 |  |  |  |  |
| Groups | T1 | T2 | T3 | T4 | T5 |
|  | Negative Control | Positive Control | Treatment Groups |  |  |
| Compound/Test item | Normal Saline | Normal Saline | Extract of *Curcuma longa* | Extract of *Ocimum sanctum* | Extract of *Ocimum sanctum* + *Curcuma longa* |
| Dosage | NA | NA | 40mg/kg daily for 14 days | 40mg/kg daily for 14 days | 40mg/kg of *Curcuma longa* and 40mg/kg of *Ocimum sanctum* daily for 14 days |
| Irradiation with Gamma rays | Nil | Irradiated with single exposure of 10 Gy of γ- radiation on 15<sup>th</sup> day | | | |

### 2.6. Gamma Irradiation

Animals of group T2, T3, T4, T5 were irradiated with 10 Gy of γ – radiation on 15th day at Indian Institute of Horticultural Research, Indian Council of Agricultural Research, Hessaraghatta Lake Post, Bengaluru-560 089 using Gamma Chamber 5000.

### 2.7. Parameters

#### 2.7.1. Growth parameter

Body weight of each group of animals were recorded on day 0,3,5,7,9,11,14,16,18 and 20 using an electric weighing balance (Indman scale corporation, India) to monitor any change in the average weight during the experimental procedure.

#### 2.7.2. Clinical signs

Cage side observation was focused on any kind of distress or physical pain that animal shown and had been recorded.

#### 2.7.3. Necropsy and Gross Pathology Examination

After the end of the treatment period (day 20), all the animals were sacrificed using Sodium thiopentone by intraperitoneal injection method (as per recommendations of CPCSEA) and subjected to gross pathological examination.

#### 2.7.3.1. Organ Weights

Testis and Epididymis weights were recorded from each animal.

#### 2.7.3.2. Tissue Collection &Histopathological Examinations

Testis and epididymis were collected from each animal and preserved in Modified Davidson’s fluid for 24 hours and were kept in 10% neutral buffered formalin (NBF) for further fixation. The fixed tissue samples were washed, dehydrated in ascending grades of acetone (70-100%), cleared in benzene and embedded in melted paraffin. Paraffin embedded samples were then cut using rotary microtome into thin ribbons of 5 micron. Routine hematoxylin and eosin staining procedure was followed to stain the slides containing tissue sections as per Luna (1968). Digital photographs of the tissue sections were taken from the stained slides using (Leica DM 2000) microscope.

### 2.8. Photography

The photography for histopathology was executed by using Leica DM 2000 compound microscope and the photos were captured by using Leica application suit; Version Lhasa application suite 4.4.0.

### 2.9. Statistical analysis

The means, standard error and standard deviation for different parameters under study was computed with the help of standard statistical procedure described by Snedecor and Cochran (1989). Tukey’s multiple range tests as modified by Kramer (1956) was used to test the difference among groups. Data were analyzed using graph pad prism version 8.0.1 software.

## 3. Results

### 3.1 Clinical signs

All treatment groups exhibited signs of radiation sickness within 24 hours after gamma radiation exposure and showed reduction in food and water intake, ru□ed hairs, piloerection, progressive decrease in body weight, abnormal breathing, hunched back, diarrhoea, pica and irritability in approach reflex. The clinical signs displayed by the animals were comparable with the findings of Koenig *et al* (2005). Irradiated animals were said to have a drastic decrease in food and water intake, weight loss over time, became lethargic, and developed diarrhoea in 2-5 days.

### 3.2 Growth parameters

#### 3.2.1 Body weight

All the groups showed gradual weight gain until the 15th day of irradiation. The negative control group and the other treatment groups had no statistically significant difference in rat body weight (P > 0.05).

After irradiation on the 15th day, however, a significant difference (P ≤ 0.0001) in body weight was seen between the irradiation positive control and treatment groups from the 16th day onwards (Fig 1). However, group T4 given *Ocimum sanctum* extract had a lower body weight loss when compared to rats given *Curcuma longa* (T3). Body weight reduction was still lower in group T5 given a combination of *Curcuma longa* and *Ocimum sanctum* than in groups T3 and T4.

**Fig 1:**
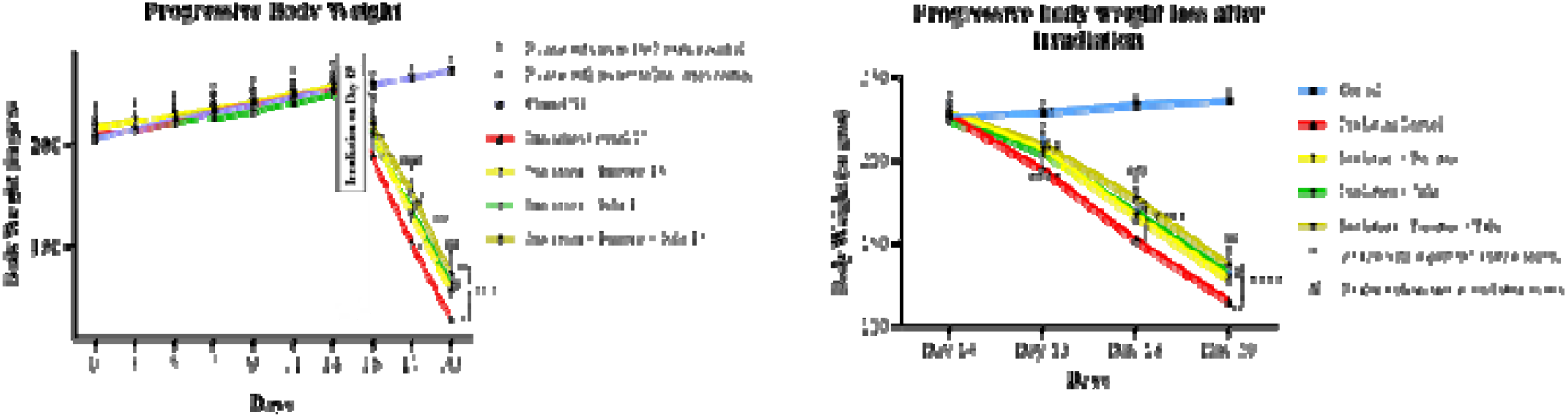
Graphical representation of variation in body weight in different time interval before and after 10 Gy of gamma irradiation.

Significant weight loss may be attributed to decreased food and water intake, loss of fluid and electrolytes due to diarrhoea and decreased absorption capacity of the gastrointestinal (GI) tract, as supported by Griffiths,(1999), Krishna and Kumar (2005), and Mihandoost *et al*., (2014) in their research.

The findings of less pronounced body weight loss in rats given *Ocimum sanctum* in groups T4 and T5 were comparable to those of Uma devi *et al*., (1999), where administration of *Ocimum sanctum* delayed the onset of diarrhoea in these animals, resulting in less dramatic weight loss with a maximum weight loss of 15-17% whereas irradiated animals treated with distilled water had a weight loss of 40%.

The less considerable body weight reduction was assumed to be due to *Ocimum sanctum’s* radioprotective action on the gastrointestinal tract. Ingestion of crude extracts of herbal remedies has been proven to decrease radiation-induced weight loss in rats, according to Jindal *et al*., (2006), Nunia *et al*., (2007), and Chaudhary *et al*., (2008).

The less substantial body weight loss in group T5 was better than in group T4, which received *Ocimum sanctum* in combination with *Curcuma longa*, indicating the possibility of a favourable synergistic effect.

#### 3.2.2. Testis and epididymides Weight

At the end of the experiment period on day 20, the testicular weights in group T2 and treatment groups (T3, T4, and T5) were significantly lower (P<0.0001) than the negative control group T1. (Fig 2). However, in treatment groups, testicular weights in groups T3 and T5 were found to be non-significantly higher than those in T4. The less decrease in testicular weights in groups T3 and T5 could be attributed to *Curcuma longa’s* radioprotective effect against testicular changes caused by gamma radiation. When comparing groups T3 and T5, the testicular weights in group T5 did not decrease as much as in group T3. This could be attributed to the synergistic effect of *Curcuma longa* and *Ocimum sanctum*.

**Fig 2:**
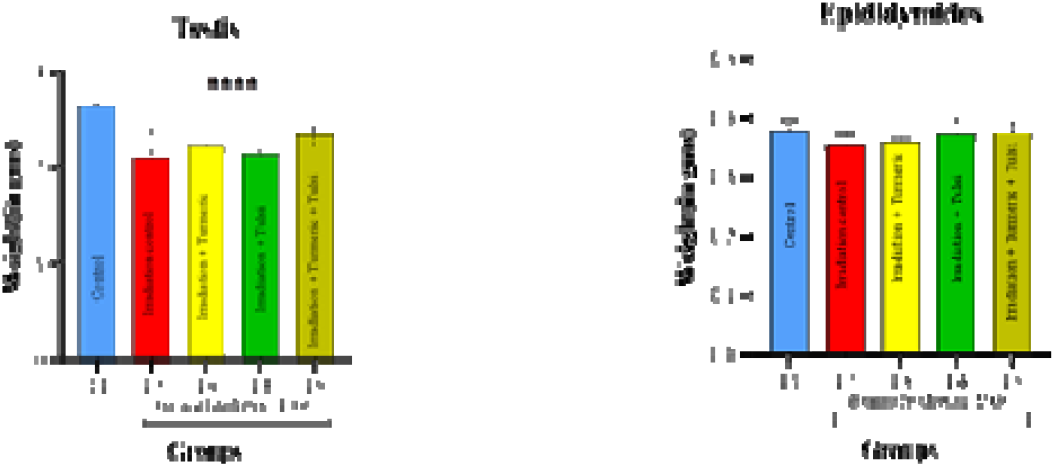
Graphical representation of variation in testis and epididymides weight in Sprague Dawley rats at necropsy.

Decreased in testicular weight after gamma radiation exposure may be due to the changes in germinal epithelial cells. (Sharma *et al*, 2011). This was further supported by (Lin *et al*.,1996) and (Jagetia *et al*.,1998) in their investigation where testicular weights and weight index declined in lethally irradiated mice.

No significant difference in epididymides was found across the groups at the end of treatment period on day 20. However, the irradiation positive control (group T2) depicted a slight decrease in the epididymal weight compared to negative control (group T1). (Fig. 2).

### 3.3 Histopathology

#### 3.3.1 Testis

The testis is one of the most radiation-sensitive tissues, and even a low dose of gamma radiation can inhibit spermatogenesis. (Ahmad and Agarwal, 2017: Micu *et al*., 2017). The different testicular cells have different levels of radiosensitivity (Beek *et al*.,1986). The mitotically active spermatogonia are the most radiosensitive in the adult rodent testis, whereas spermatocytes and spermatids are more resistant to ionizing radiation. (Aitken and Iuliis, 2010).

Light microscopy of cross-sections from the negative control (group T1) revealed normal testicular architecture, seminiferous tubule integrity, and interstitial tissue with Leydig cells found in groups. The seminiferous tubules showed normal spermatogenesis, with normal morphology of all germ cell lines including spermatogonia, primary spermatocytes, secondary spermatocytes, spermatids, and spermatozoa. (Fig 3: A&B).

**Fig 3:**
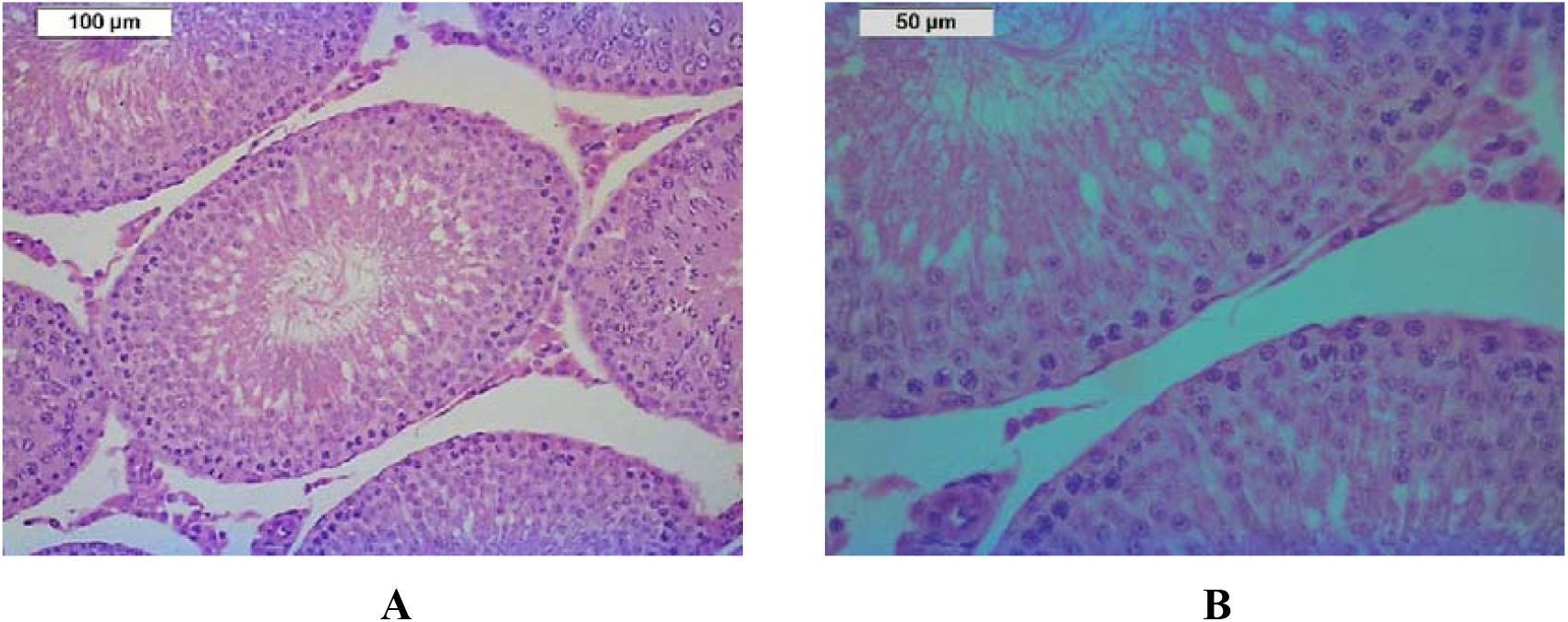
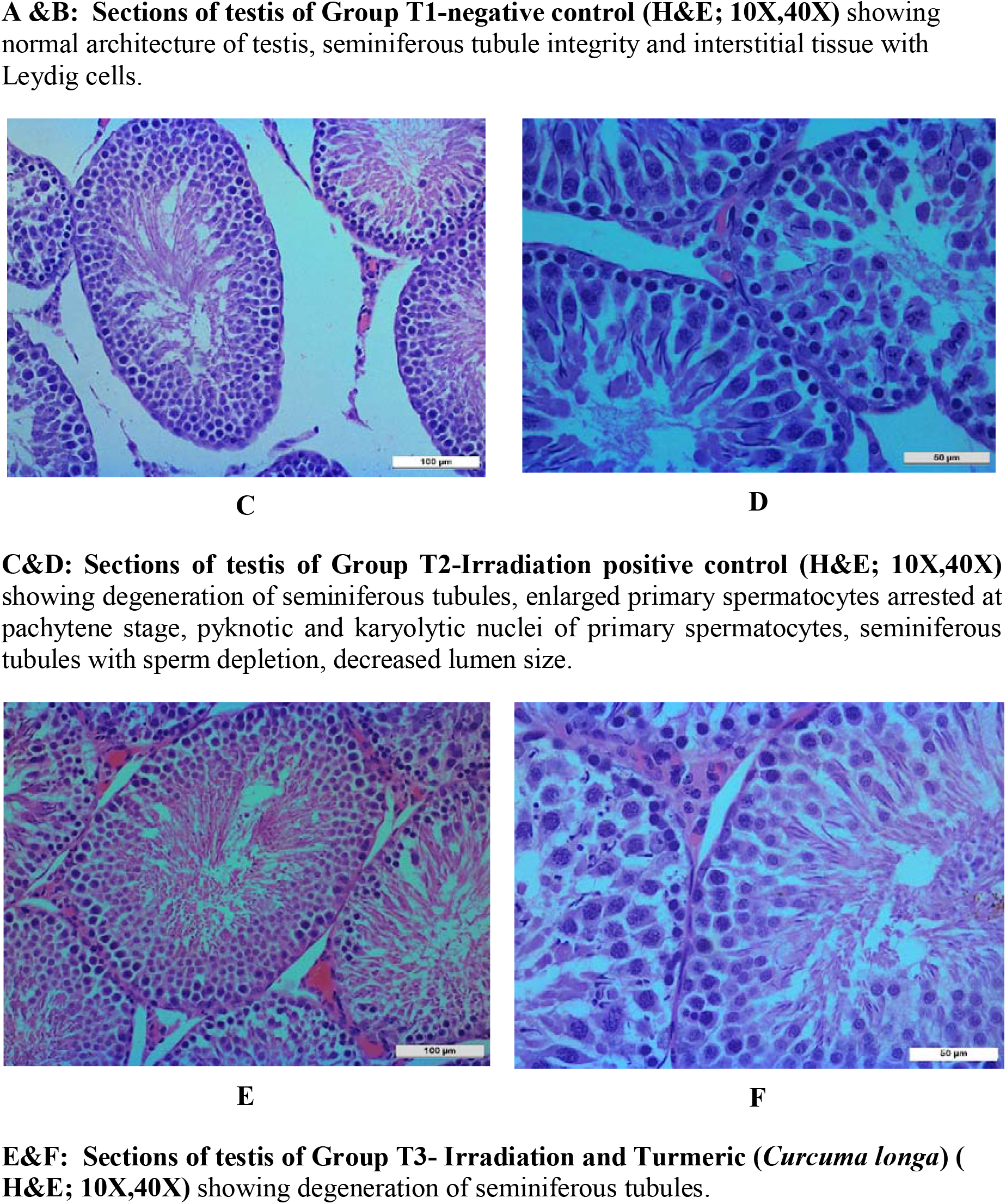
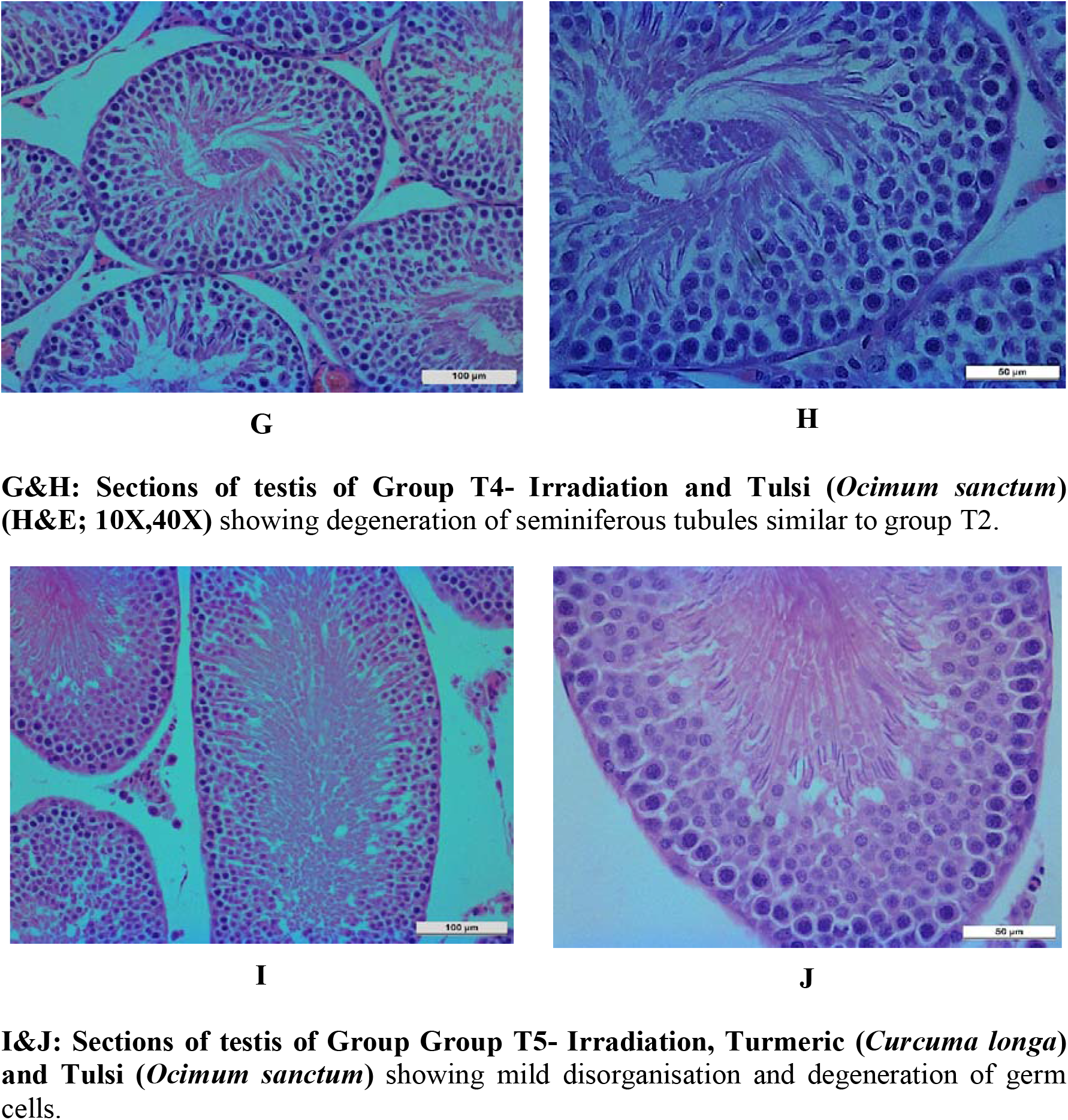
Histological stained sections of testis in different groups.

In Group T2-Irradiation positive control, microscopic images showed the seminiferous tubules were degenerated. The seminiferous tubules were disorganized and the basement membrane was irregular. A significant decrease in diameter of the seminiferous tubules was observed. Spermatogenesis was found to be disrupted in most of the tubules. Primary spermatocytes showed a high level of responsiveness. The nuclei of the majority of primary spermatocytes were found to be enlarged and deeply stained, some having pyknotic nuclei. Secondary spermatocytes were found to be less affected compared to primary spermatocytes. Also, spermatids showed a low level of reactive changes. However, decreased spermatids density was found in lumen of the seminiferous tubules. The Leydig cells in the interstitial tissue had pyknotic nuclei. There was presence of some unusual spaces and dilations associated with lymphocytes and plasma, which indicated inflammation in the interstitial tissue. (Fig 3: C&D).

In Group T3, microscopic images showed degeneration of seminiferous tubules, arrest of spermatogenesis. Primary spermatocytes were significantly affected which were comparable to irradiation positive control (group T2). The spermatids density was not much affected. Disorganization of germ cells was also observed. (Fig 3: E&F).

In Group T4, microscopic images showed degeneration of seminiferous tubules. Enlargement of primary spermatocytes and pyknosis of nucleus was seen but they were comparatively less than irradiation positive control (group T2). Disorganization of germ cells, depletion of germ cells was observed (Fig 3: G&H).

In Group T5, microscopic images showed mild degeneration of seminiferous tubules, disorganization of primary spermatocytes were less comparatively less. Spermatids density was not much affected and appeared almost normal to group T1. (Fig 3: I&J).

#### 3.3.2. Epididymis

By light microscopy of the cross sections. Group T1-negative control showed normal epididymal ducts with columnar epithelia and stereocilia. The epithelium was surrounded by an intact basement membrane made up of basal cells. There was an abundance of spermatozoa in the lumina of the epididymal duct. (Fig 4: A&B).

**Fig 4:**
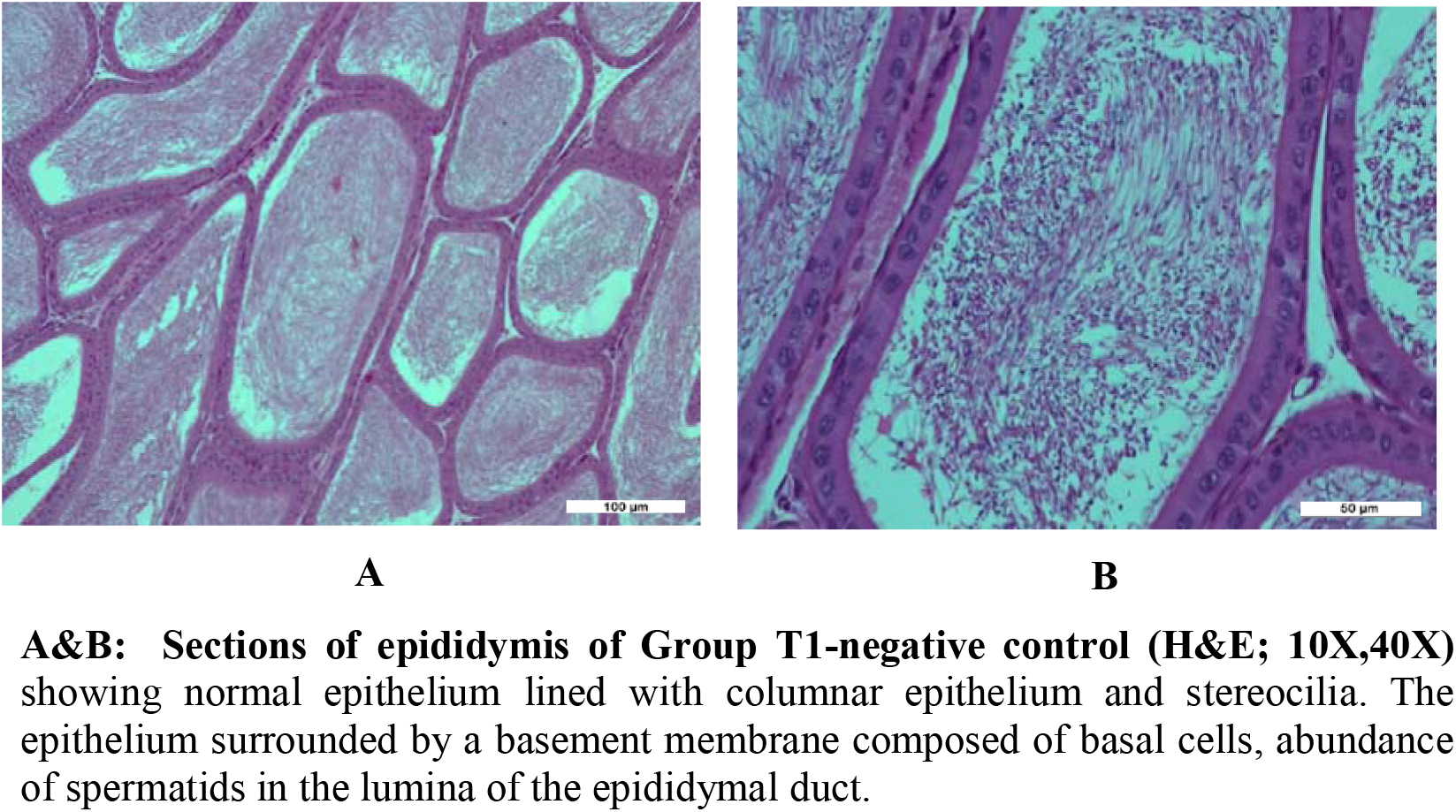
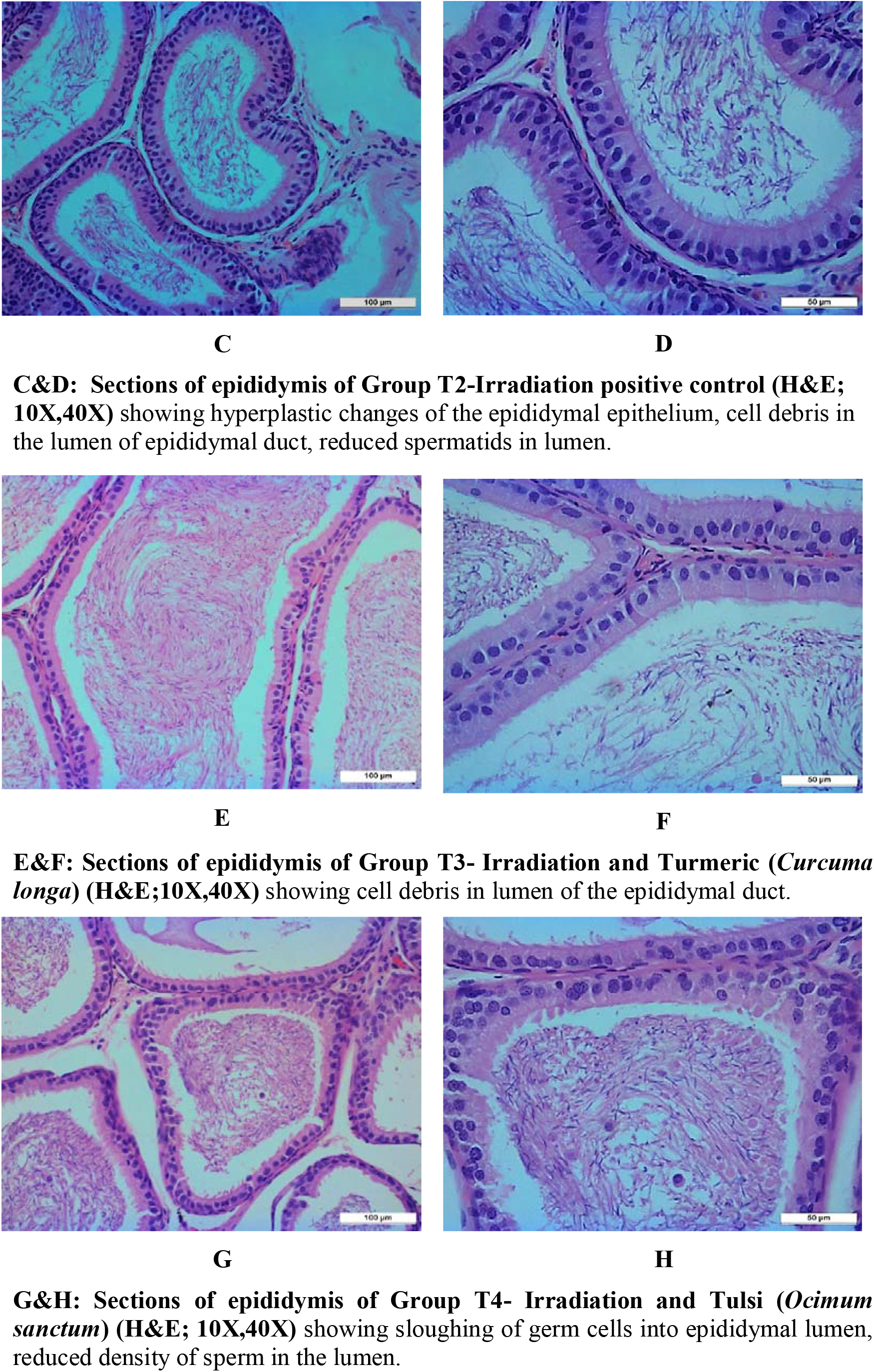
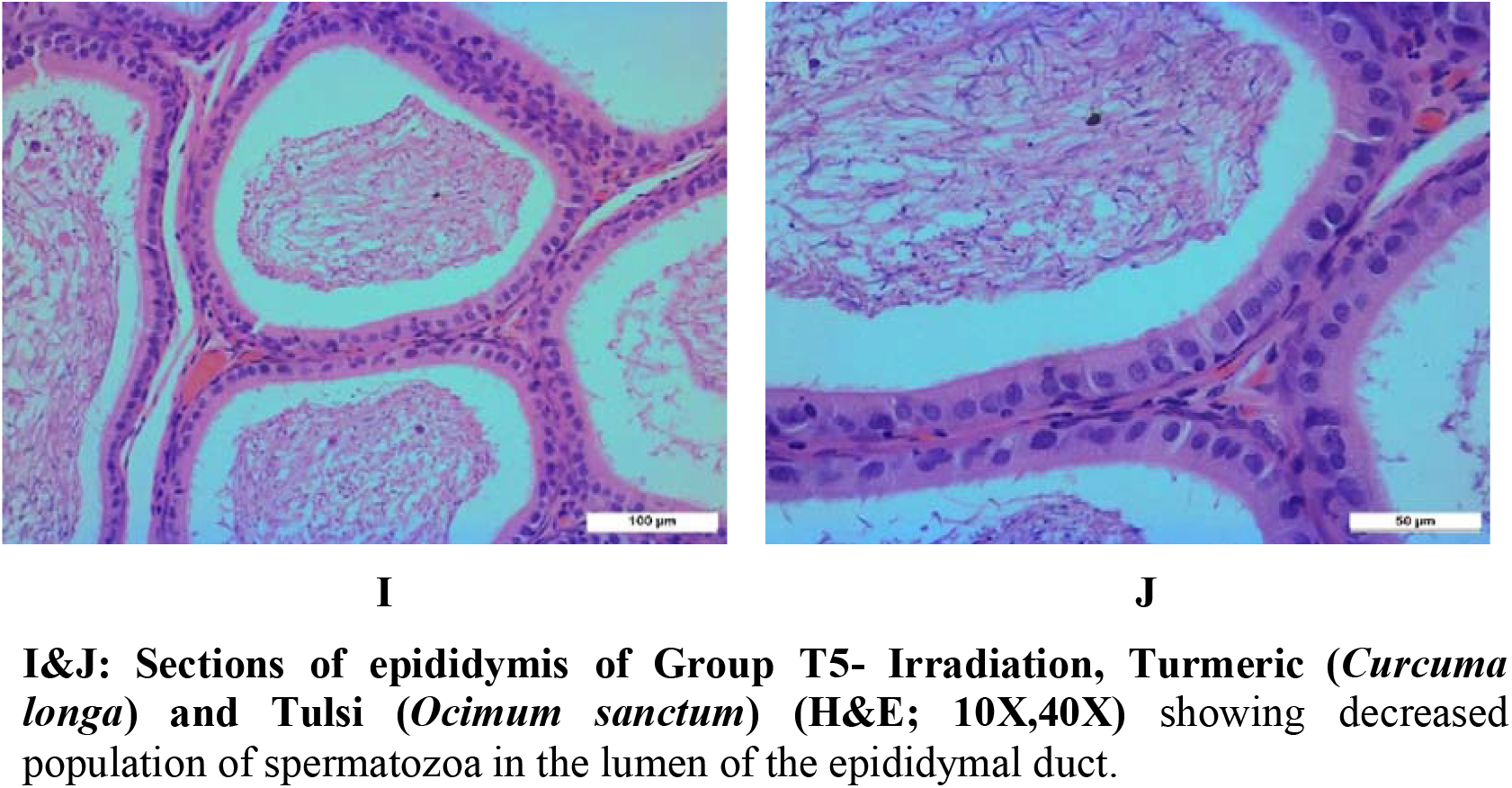
Histological stained sections of epididymis in different groups.

In group T2, microscopic images of the epididymal epithelium revealed hyperplastic changes in the epididymal epithelium. The nucleus of the epididymal epithelium was found to be elongated. To varying degrees, the epididymal tissue was distorted. There was cell debris in the epididymal duct lumen. There was a significant decrease in spermatid count in the lumen compared to negative control group T1. (Fig 4: C&D).

In group T3, the spermatid content in epididymis lumen was reduced. There was presence of cell debris in the lumen of epididymal duct (Fig 4: E&F).

In Group T4, microscopic images similar hyperplastic changes in the epididymal epithelium similar to irradiation positive control (group T2). Sloughing of germ cells into the epididymal lumen was observed. There was presence of cell debris in the ducts, reduced density of spermatids in the lumen of the epididymal duct. (Fig 4: G&H).

For Group T5, microscopic images demonstrated cell debris in the lumen of epididymal duct and decreased population of spermatids in the lumen of the epididymal duct (Fig 4: I&J).

## 4. Discussion

The present study shows that single dose gamma irradiation of 10 Gy, induced histopathological and morphometric changes in testis in irradiation positive control (group T2) and all treatment groups (T3, T4 and T5). The histopathological changes of testis in the irradiated rats were comparable to Sharma *et al*., (2011) in which the peripheral and central tubular diameter of seminiferous tubules decreased progressively after irradiation, possibly due to spermatogenic cell loss and tubular disorganisation. This was consistent with the findings of Kumar *et al*., (2007); Samanta, and Goel, (2002) in which animals that had been irradiated had shrunken and empty seminiferous tubules that were linked to a decrease in total germ cell population.

The high level of radiosensitivity of spermatogonia to gamma radiation observed in our experiment were comparable to studies carried out by Mian *et al*., (1977) in which the altering spermatogonia were more sensitive to ionising radiation and were destroyed by induction of a dosage close to 3 Gy/h in Sprague-Dawley rats and mice. Similar pattern of alterations were observed by Dewey *et al*. (1991); Chen *et al*., (2000); Seong *et al*., (2001); Cordelli *et al*., (2003) in which they have reported testicular germ cells as the most radiation-sensitive cells. These findings can further supported by studies carried out by (Hussein *et al*., 2006) where spermatogonia were found to be the most vulnerable cell types, and their numbers rapidly decrease after exposure to ionising radiation. Only a small percentage of mature germ cells and testicular stem cells survived. While mature germ cells were lost during maturation, the process of stem cell recolonization begins.

Agarwal *et al*., (2012) reported that the appearance of significant spaces in the germinal epithelia and disruption of the spermatogenic series observed in some of the treatment groups, indicating structural disorganization, which indicated testicular cytotoxicity or necrosis were consistent with our present results. Boorman *et al*., (1999) reported that a drop in sperm content was expected when spermatogenic disruption occurs in the testis.

Gamma irradiation causing histopathological changes in testis observed in this experiment was comparable to studies carried out by Pareek *et al*., (2005) where gamma rays cause degenerative effect on spermatogenesis in lethally irradiation mice. These findings were further supported by Peltova *et al*., (1992) in which alterations to normal testicular architecture was triggered by suppress in its dual character steroidogenic, and spermatogenic activity. Radiation further induces oxidative stress, suppressing antioxidant mechanisms, and activating numerous molecular pathways involved in germ cell life and death. These findings were consistent with studies carried out by Sapp *et al*. (1992); Zhang *et al*., (2001) in which the loss of germ cells after ionising radiation exposure was linked to apoptosis. Similar pattern of alterations was observed by Agarwal *et al*., (2014) where gamma radiation impairs Leydig cell activity, resulting in spermatogenesis disruption, spermatozoa depletion, and testicular atrophy by creating reactive oxygen species (ROS). These findings were consistent with Kerr *et al*., (1993); Bakalska *et al*., (2004); Yang *et al*., (2006) where degeneration (pyknotic nuclei) or death of spermatocytes and spermatids depicted that meiosis and spermiogenesis had been disturbed.

Histopathological observations in the present study clearly depicted that pre-treatment with combination of *Curcuma longa* and *Ocimum sanctum* in group T5 prior to irradiation exposure had synergistic effect and appreciably showed radioprotection on the histology of testis. At the same time, group T3 and group T4 given an extract of *Curcuma longa* and *Ocimum sanctum* respectively alone prior to irradiation showed minimal noticeable radioprotection on histology of testis.

Study found that *Curcuma longa* not only has a non-toxic effect, but it also has cytoprotective effects on the histoarchitecture of the testes in diabetic rats. (Olanrewaju *et al*., 2017). Rashid and Sil, (2015) findings confirmed the protective effect of curcumin and indicated that treatment of diabetic rats for 8 weeks reduced testicular alterations and gave optimum protection at a dose of 100 mg/kg body weight, indicating that curcumin protection against oxidative stress-mediated testicular damage. Earlier findings by Sudjarwo *et al*., (2017) depicted that Curcumin’s had preventive impact on lead acetate-induced reprotoxicity (50 mg/kg BW), discovered that daily oral administration of curcumin at three doses (100, 200, and 400 mg/kg) to rats for 40 days improved the histological structure of testis.

The current study found that 10 Gy gamma irradiation caused histopathological changes in the epididymides of irradiated animals of irradiation positive control and other treatment groups.

Exfoliation and sloughing of testicular germ cells into the epididymal lumen occurs either as part of the maturation process (peripubertal animals) or secondary to injury and/or sloughing of the testicular germ cells. The lower sperm density in the epididymis of the irradiated animals may be attributed to reduced spermatogenesis in the testis due to multiple injured seminiferous tubules reported by Nna *et al*., (2017) was comparable to our findings.

Sloughing of germ cells into the epididymal lumen observed in Group T4-(Irradiation and *Ocimum sanctum*) can be further supported by the findings by (Foley, 2001) in which an increase in the number of sloughed, degenerate germ cells in the epididymal lumen often indicates continued testicular degeneration/atrophy, spermatid retention, or germ cell exfoliation in adult rodents.

Histopathological observations of epididymis in the present study clearly depicted that pre-treatment with *Curcuma longa* in group T3 appreciably showed moderate degree of radioprotection in epididymis after exposure to gamma radiation with less affected spermatids density in the lumen of epididymal duct. While the group T4, pre-treated with *Ocimum sanctum* demonstrated not much radioprotection in epididymis of rats. At the same time, group T5 pretreated with extract of both *Curcuma longa* and *Ocimum sanctum* did not show much radioprotective histopathological differences.

## 5. Conclusion

Exposure of animals to 10 Gy gamma radiation resulted into significant decrease in body weight and testicular weight. Histopathological observations revealed testicular degenerative changes in the seminiferous tubules with primary spermatocytes being most affected, enlarged primary spermatocytes arrested at pachytene stage, pyknotic and karyolytic nuclei of primary spermatocytes, seminiferous tubules with sperm depletion, decreased lumen size. Epididymis showed hyperplastic changes of the epididymal epithelium, cell debris in the lumen of epididymal duct, reduced spermatids in lumen.

